# The Study on Computer Vision-Assisted Cell Bank Construction and Screening & Classification

**DOI:** 10.1101/771089

**Authors:** Feng Yanmin, Chen Hanlong, Bai Xue, Chen Yuanyuan, Dai Yuguo, Feng Lin

**Affiliations:** School of Mechanical Engineering & Automation, Beihang University, Beijing 100191, China; Beijing Advanced Innovation Center for Biomedical Engineering, Bei hang University, Beijing 100083, China

**Keywords:** cell factory, computer vision, cell segmentation, recognition

## Abstract

Computer vision technology plays an important role in screening and culturing cells. This paper proposes a method to construct a helper cell library based on cell image segmentation and screening. Firstly, cell culture and image acquisition were carried out. The main content is to use laboratory conditions to carry out different cell types. Through careful observation of the whole process of cell proliferation and passage, the representative pictures of different stages were taken. Analysis and summary of the relevant morphology, texture, color characteristics. Secondly, computer vision technology is used to segment cells and extract the main features such as cell perimeter and area. Explore the automatic information extraction method of cell bank, and complete the image segmentation of individual cell image from the whole picture. Finally, the cells were screened and identified. Investigate different pattern recognition methods and neural network structures, and prepare pictures of various cell pictures. The corresponding neural network and prediction program are constructed. This paper proposes an automatic image processing method for each image category in cell culture cycle, which improves the automation of production process. At the same time, compared with the design of a single algorithm for a certain type of cell, different algorithm design ideas are proposed for three types of pictures with different characteristics, which is closer to the dynamic change of cell morphology in the process of cell culture. This research has important application prospects in promoting cell factory research, cell bank construction and automatic screening.

## I Introduction

Cell factory is a recently developed mass cell culture technology. The utility model has the advantages of good aseptic operation, small operation space and high degree of automation. Cell factory is widely used in vaccine manufacturing, cell engineering and monoclonal antibody manufacturing industry(Aliperta et al., 2017; Barzegari et al., 2014; Forbes et al., 2015; Hong and Nielsen, 2012). In our lab, we have developed single cell operation (Feng et al., 2016; Feng et al., 2017a; Feng et al., 2013; Feng et al., 2017b). However, animal cells are diverse and require different culture environments(Feng et al., 2018), which poses a major challenge to cell culture in vitro. Because large-scale cell culture needs to summarize the mode of scale and standardization, but there are differences in the culture environment of cells(Fan et al., 2017). So, it is difficult to carry out large-scale mass production on the basis of traditional artificial culture. At present, 96-well plate and 384-well plate are widely used in the field of biomedicine to cultivate a large number of cells(Sykes and Avery, 2009). This method requires researchers to observe orifice plates with a microscope. The whole process takes a long time and requires skilled staff. Automated Cell Screening Platform is an automated operating system developed rapidly in recent years, which uses visual information to complete a series of automated training steps (Breker et al., 2013). The cell screening system based on computer vision technology can realize image acquisition and processing, which is the key part of the automatic cell screening platform (Roy et al., 2002). With the help of the automatic cell screening platform, the experimenter can observe the cell image in real time through the biomicroscopy (Roy et al., 2002), and then stitch, segment, extract and count the cell image. Then, the functions of feeding and moving can be accomplished by controlling the driver to screen the cells in the image. The whole process requires less people to participate in the link, improves work efficiency and reduces staff errors caused by long hours of work, which greatly improved the accuracy of screening and overall reduced economic and human costs. Therefore, it has great potential for development to use of computer vision for automatic cell screening(Sommer and Gerlich, 2013).

In addition to its application in large-scale cultured cells, computer vision technology plays an increasingly important role in biomedical fields such as pathological identification of cells (Anthimopoulos et al., 2016; Mohamed et al., 2018; Tourassi et al., 2016). Automated testing is an urgent need of modern hospitals. At present, the identification of cells is mainly accomplished by manual work, and the results of identification largely depend on the accumulated experience, which has a high demand on practitioners and a strong influence on subjective factors. In order to improve the accuracy, long-term training is needed, and the technical ability and mental state of operators are highly demanded. Many factors restrict cell identification.

In 2018, a large number of melanoma cases were diagnosed in the United States. Nearly 10,000 people are expected to die from melanoma(Saraswathi, 2017). Demand for systems with high accuracy and automatic detection of skin cancer-related conditions is increasing. Computer software can automatically detect and analyze areas related to skin cancer from captured digital images, and patients can test at any time. Corresponding inspection procedures have been developed and more than 90% accuracy has been achieved(Saraswathi, 2017).

With the development of computer vision technology, image segmentation and other algorithms can separate individual cell images from image samples under a microscope, that is, only a single cell exists in the obtained image. On this basis, it is convenient to extract different features of cells-color, shape and texture. Then we further summarize the characteristics of higher level, such as cell area, perimeter and so on. Traditional pattern recognition methods such as back propagation network and SVM use the above features as input to complete cell classification, so as to achieve the purpose of cell classification. But when it is really used in life and production, the sampled pictures have different characteristics with the sampling conditions and disease types. A series of algorithms including pre-processing, segmentation and feature extraction are designed for each specific project(Boucher et al., 1998; Moen et al., 2019; Van Valen et al., 2016). The level of the designers and the selection of parameters in the algorithm can easily affect cell recognition. In this way, only a part of the characteristics summarized by experience are extracted in the process of manual participation, which leads to a large amount of waste of information, so it has limitations. Therefore, it is usually necessary to cooperate with manual aided recognition after automatic machine recognition.

Compared with foreign countries, domestic research on automated cell screening platform is less. Nowadays, cell screening and culture are facing many problems, such as too long detection time, tedious selection process and so on. And foreign countries have a leading position in automatic cell screening system. Molecular Devices has designed and manufactured ClonePix, the world’s first fully automated cell screening system(Agarwal et al., 2017), which improved the traditional imaging screening method to screen suitable cells by fluorescence imaging. AVISO Germany developed a fully automatic cell screening platform. This system has the functions of cell recognition, cell screening and automatic generation of biology important information report forms. At present, there is no mature market-oriented automatic cell screening system successfully developed in China.

Although there are many ways to segment cell images (Al-Kofahi et al., 2018; Anoraganingrum, 1999), it is necessary to track the cultured cells until the cell completes its periodic proliferation in order to study the effective image processing and segmentation method of cell factory in every stage of cell culture. We should take pictures at each representative stage to get the cell image, and then design the corresponding Algorithms method according to local conditions. Cells show different characteristics with the time of culture. There is a problem of insufficient generality in individual identification methods. Therefore, sufficient and representative images should be selected from the whole process of cell proliferation and passage to assist the algorithm design and improvement. Based on the different stages and different kinds of cell images, the different visual characteristics are analyzed, summarized and categorized, and then different image preprocessing and segmentation algorithms are compared and selected. According to the different features of the cell image after classification, the corresponding segmentation processing scheme is designed, and then the program is compiled, and the parameters are updated according to the test results to complete further optimization and debugging. According to the different morphological characteristics of different kinds of cells, the classification algorithm is designed and the program is developed and debugged. The network structure and parameters are adjusted continuously according to the feedback so that the training network can converge successfully and identify different kinds of cells successfully.

There are many studies on the recognition and classification of blood cells by vaccines. aRelevant human health information can be obtained by automatically identifying red blood cells and obtaining statistical data (Nie et al., 2017).

In this paper, the morphological characteristics of cells under the microscope are divided into three categories. By using Canny operator, Hough detection, morphological operations such as expansion, corrosion and watershed, edge detection and other cell segmentation methods, the image of the cell region is segmented from the image, and the cell count and feature extraction are realized. Finally, the recognition between different cells is completed.

## II Cell culture and Image Acquisition

### (1) Characteristic Analysis of Cell Image

Different cells were cultured under laboratory conditions, including cancer cells DU145, MCF-7 and immune cell RAW246.7. And the whole process of cell proliferation and cell passage operation was carefully observed (Fig. 1A). We realized the standard operation of cell passage, resuscitation and culture in the laboratory. Representative pictures of cell different stages were taken using microscopy (Fig. 1B). Through analyzing and summaring of the relevant cell morphology, texture, color characteristics, we concluded the feasibility of cell factory and the necessity of real-time cell monitoring. The cells should be passaged when they reach about 90 % confluence. Because adherent cells can produce contact inhibition, which is an important characteristic for cell proliferation, so we should consider the culture area as a threshold, using visual recognition to detect the cell growth area and automate the feeding and replacement operation in the large-scale cell culture. In the experiment, RAW264.7, MCF-7 and DU145 were subcultured for two to three times, and a large number of cell images were obtained. The characteristics of freshly dissociated cells in short-term culture was as follows:

**Fig. 1.**
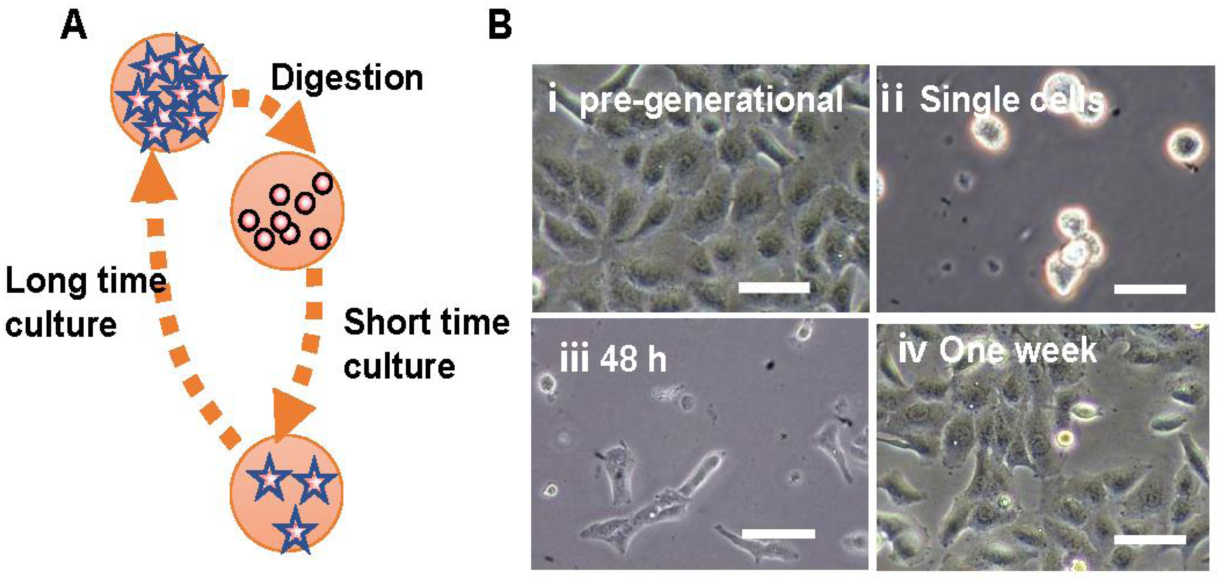
Different cells exhibit different physiological characteristics at different growth stages and need to be classified and studied. (A) the schematic of cell culture operation process. (B) Cell morphology represented by DU145 cells at pre-generational (i), digested into single cells (ii), cultured for 48 h (iii), and one week (iv). Bar, 50 μm.

Cells will be dissociated from the surface of the Petri dish after trypsin digestion and be suspended in to single cells in culture medium after spire up and down. Then the single cell suspended in culture medium was round, some cells overlapped, furthermore cell contour imaging was clear (Fig. 1B ii).

It was found that the transmittance of the cytoplasmic part was higher after the overall observation of the photographs taken. it could be better distinguished from the nucleus and cytoplasm for some cells, but some cells had darker imaging and stacked with other cells without obvious and clear boundary lines. The nucleus is difficult to be clearly distinguished from cytoplasm. In addition, RAW264.7 cells are also round when suspended in cell culture medium, which is not different from cancer cells after treated by trypsin in visual characteristics. Therefore, the algorithm is designed together.

After dissociation, the cells suspended in the culture medium were spherical. After adhere to the Petri dish, the cells will gradually extend into flat rather that round in cell culture medium and tightly adhered to the growth surface of cell culture substrate. At this point, when viewed under a microscope, the cells present a flat, bright circle. After that, the cells began to proliferate, showing irregular patterns. At this time, the cell outline is irregular, but the cell outline is clear, and there is a clear distinction between the dark background. Therefore, it is easier to distinguish cells by the changes of light and shade at the cell boundary, as shown in the Fig. 1B iii. With the increase of culture time, cells begin to connect tightly. The junction of cells is brighter, while the nucleus of cytoplasm cells is darker. It is impossible to observe the clear nucleus with the human eye, which makes the extraction of the components of each part of the cell very complicated (Fig. 1B iv). MCF-7 cells are more typical, so the analysis is based on this type of cells, while the cell outline of DU145 is irregular and the cell boundary becomes difficult to distinguish.

### (2) Design of Cell Preprocessing and Segmentation Algorithms

According to the corresponding algorithm investigation, there is no standard and unified cell extraction process. Generally, we should design image preprocessing according to local conditions. For strongly adherent cells, relevant studies(Mukherjee et al., 2012) used Hough transform circle detection to separate images, which uses the color gradient method to set the threshold and segment the edges of cell adhesion, and then extracts the features. This method is different from the gray threshold detection. However, we find that in MCF-7 cells show a variety of morphologies with different size and shape, At the same time, K-means, as a common and classical color classification algorithm, can classify multiple samples into different categories in multi-dimensional image space. In this paper, it is observed that there are obvious differences in color brightness between cell boundaries and cell parts of dense adherent cells, which makes it possible to use K-means to cluster cell images. In this paper, we propose to use edge detection and circular detection as the core to detect and count the circular cells represented by RAW264.7 cell lines. It is mainly from the following aspects to brush for adherent and irregular cells: one is color feature brushing, which is used to verify whether the clustering algorithm can distinguish the boundary from the interior of the cell. The other is that practicability of watershed under specific conditions is determined by experiments, and image enhancement is carried out by various cell image preprocessing methods to improve the recognition accuracy.

Above all, three kinds of cell conditions were experimented with the above algorithms, and the process was further optimized after summarizing the inductive methods in the experiment following the technological process as showed in Fig. 2.

**Fig. 2.**
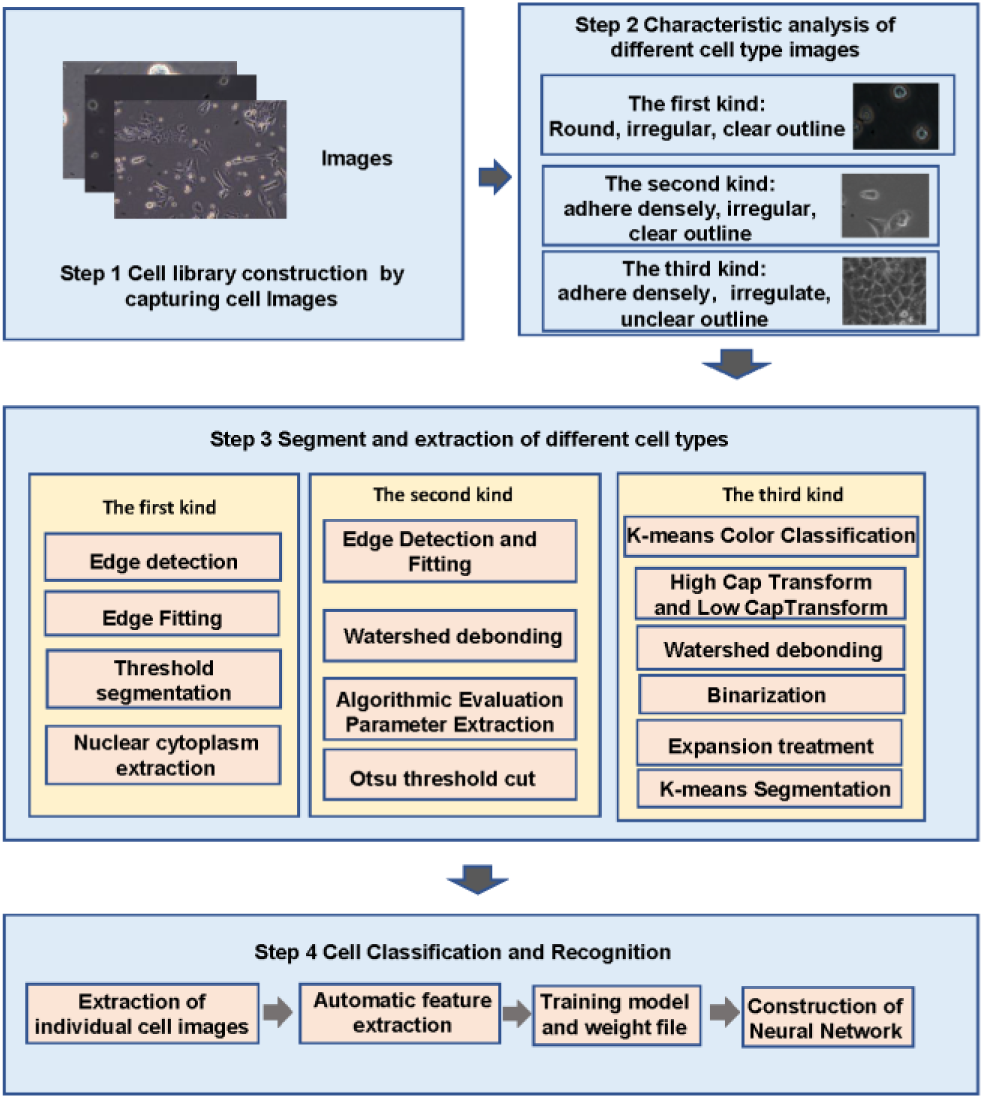
Overview map of three different types of cell segmentation and recognition, including four steps: different types of cell library construction, characteristic analysis, segment and extraction and cell classification and recognition.

## III Cell Screening and Recognition based on three cell morphological characteristics

Cell image processing technology involves many basic steps, first of all, image acquisition, then image preprocessing, and subsequent counting and extraction operations. The successful completion of image segmentation lays a solid foundation for image post-processing and data acquisition. In this section, we will discuss in detail the effect of different algorithms on image segmentation under different cell morphological characteristics and the corresponding steps.

### Technology used in this paper

OpenCV is one of the most widely used libraries in the field of computer vision. It can be compatible with various systems and environments and has good cross-platform performance. OpenCV contains a large number of image processing methods, so it is widely used to solve various image processing problems. OpenCV encapsulates a variety of perfect visual processing related functions and functions, and leaves a wealth of interfaces to facilitate diversified image processing. This topic installs OpenCV Library in Python environment, and then calls library functions to perform various complex image processing. In the follow-up experiments, the main task is to capture a large number of cell images. In the process of developing the program after the algorithm design, the image can be preprocessed, segmented and extracted easily with the help of OpenCV.

Jupyter Notebook is a Web application that focuses on interactive computing and has developed rapidly in the field of scientific computing. It can be applied in various fields such as developing programs, generating documentation, code execution, and displaying strong interactive results. Jupyter Notebook allows users to create and execute code directly on the Web. The results of code execution are also displayed directly below the code block, which helps users debug at any time. And the code block can flexibly change position. You can write multiple code blocks and explanatory files and comments directly on the same page. This is very useful for software development and interpretation.

This paper uses Anaconda to solve the installation problems of Jupyter Notebook and other modules. Anaconda has automatically installed Jupyter Notebook and some other tools, including a variety of commonly used scientific computing packages supported by Python. Anaconda can be easily used to configure and share the environment. The cloud synchronization environment supported by Anaconda can solve the problem of different configuration files installed on different platforms to a certain extent. It has many advantages of choosing Python to conduct our topic. The development of GUI can be easily combined with OpenCV. When developing with C++, the use will involve different compilation environment problems. In order to integrate OpenCV with Qt development, developers need to compile themselves for subsequent development. For example, Qt5 compiler based on MinGW53 0 takes more than 30 minutes to compile. Python supports the embedding of GUI development tools such as wxPython and PyQt. It only needs to install plug-ins in compiler environment such as PyCharm. Python has a complete OpenCV library, which reduces the time cost.

The working environment of this paper is version 1.9.7 of Anaconda. In this project, a separate environment is installed, which is different from the root environment, and a virtual environment is created to manage each package. OpenCV can be installed by PIP instructions. The environment of this paper is OpenCV 4.0.0. Then the processing functions and programs based on this are written, and only the necessary packages are installed when needed, which can reduce the existence of redundant packages and facilitate the subsequent upload and download of cloud environment.

### The first kind of cell image: cell edge detection and circle fitting using OpenCV

#### Design of cell extraction algorithms

To recognize and extract cells, edge detection is needed, and then the detected edges are fitted. If there is a big difference between the cell and the background color, the target (cell) and background can be segmented by threshold, which segmentation can be used to isolate target (cell) and background. Common methods include Otsu segmentation, binary segmentation, etc. Otsu segmentation is an adaptive threshold method without manual setting. So it is widely used in all kinds of segmentation(Pavesic and Ribaric, 2000). In this method, the greater the difference between background and target, the better the segmentation effect. Otsu input gray-level features, according to the different features of background and target in gray-level image, the image is divided into two categories and the binarization is completed at the same time.

In the algorithm, a threshold T is set as the criterion of gray dichotomy. The ratio of the number of pixels judged to be the target (cell) after dichotomy to the number of all pixels in the picture is marked as ω_0_. The average gray scale of these points is recorded as µ_0_. The proportion of the number of non-target pixels to the total number of pixels is ω_1_. Its average gray level is µ_0_. µ is the total average gray level of the whole image. When the variance g between the average gray level of the two classes and the average gray level of the whole pattern is the largest, it means that the error classification is the smallest, and then the segmentation is completed.

The image size was set as X×Y. When the background color is darker (that is the actual situation of this experiment), the number of pixels whose gray level is less than T is denoted as N_0_, and the number whose gray level is greater than T is denoted as N_1_.

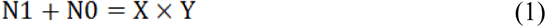

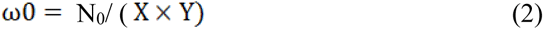

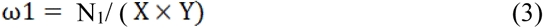

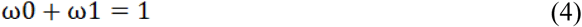

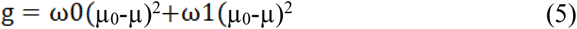

OTSU method was applied to judge the specific situation in the experiment. From the image analysis, some cells are judged as part of the background because of the small difference between the color and background of the central part of the target. The cells in the picture are not all in the same focal plane because of their suspension state. Among them, some cells have darker imaging. Otsu will cause great loss in processing images. The most important thing is that the central part of some darker cells is recognized as the background (Fig. 3D), resulting in only the edge of the break, which phenomenon results in a great loss of information. Other methods such as threshold selection in binary segmentation have strong randomness, which will bring great inconvenience to single threshold segmentation. If the threshold is too high, a lot of information will be lost and the complete image will not be obtained. While if the threshold selection is very low, background noise or impurities will be detected, which is not universal, so other ways should be considered. In the picture, the edge part of the cell is obviously distinguished from the background, so we consider using the edge information of the cell to extract the cell. Nowadays, there are many methods of edge detection. OpenCV has its own find Counter function, but it requires a high degree of edge integrity. So we consider other ways to extract edges, among which Canny edge detection algorithm is very effective(JOHN CANNY, 1986). The edge detection of Canny algorithm can be divided into four steps: the first one is Gaussian Smooth filtering, the second one is that differential convolution Kernel Enhanced Image Edge. The third one is suppression of non-maxima. The fourth is double Threshold Segmentation: setting one or two thresholds T1 and T2. If the edge gradient of some point is greater than T1, the point is considered to be an accurate and reliable edge point, which reduces the probability of error detection of non-edge. However, due to the high threshold setting, the resulting image edges are less, and it is difficult to be fitted in the follow-up work, so another low threshold T2 is set. After high threshold image processing, the edges are linked into contours. When the endpoints of the contours are reached, the algorithm searches for the points satisfying the T2 restriction in the neighborhood points of the breakpoints, and finally finishes the acquisition of the points on the edges.

**Fig. 3.**
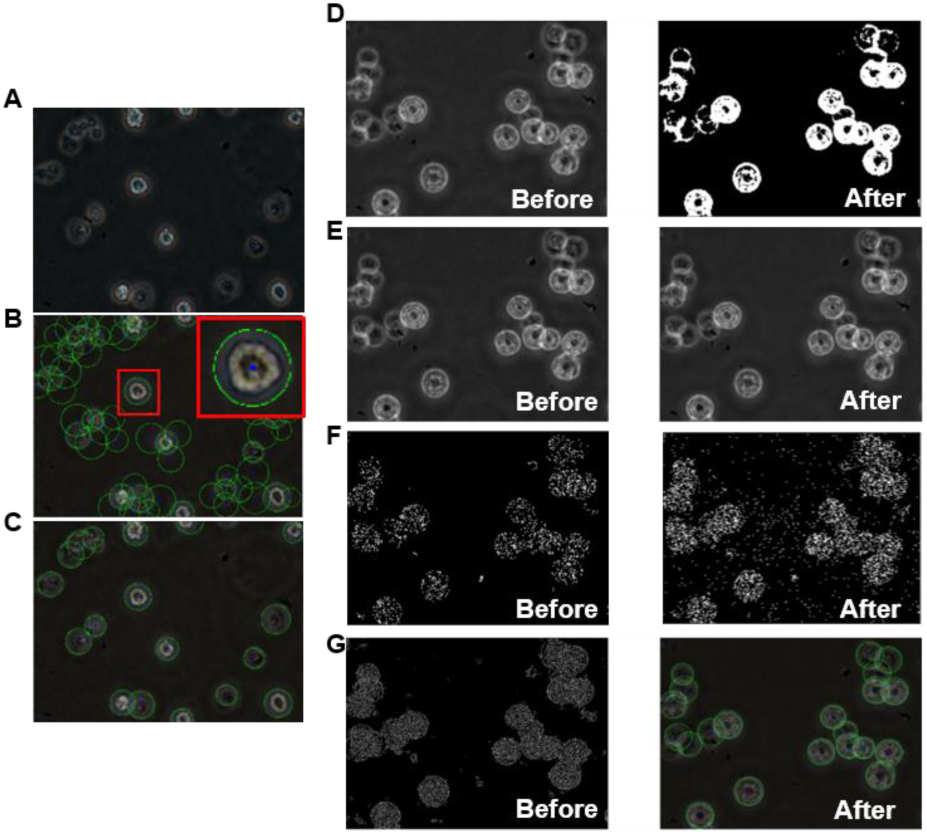
Circle-like cell segmentation and recognition. The binary image of background noise is filtered out, then the edge contour is detected and extracted. Finally, the cell edge is fitted by circle and counted. (A) original image circle like cell.(B) Image before debugging in the process of cell circle feature extraction and debugging. (C) Image after debugging in the process of cell circle feature extraction and debugging. (D) images of before and after Otsu treated. (E) Image before and after median filtering. Comparisons of unused pictures with median filtering(F) Canny image before and after median filtering. Canny operations were compared using median filtering and unused pictures.. (G) Hough detection effect.

At the same time, there is mechanical and electronic interference in the microscopy of the experiment, and the signal will be disturbed in the transmission process. There is some noise in the cell picture taken. Due to the movement of camera during observation and the flow of cell culture medium during photography, the movement of collected cells will be blurred. Median filtering method can eliminate noise and reduce interference before edge detection. Median filtering is a kind of non-linear digital filtering, which can reduce the influence of outliers without reducing image contrast (Fig. 3E). When processing gray image with median filter, the median of each point covered by the image core is used to replace the gray value of the central point of the detection area, so traversing. We find that the image of the function is enhanced, which greatly improves the effect of Canny operation.

Because of the high resolution of the microscope, the small noise is not obvious. But the difference is obvious after Canny processing. It can be seen that the result of Canny operation after median processing has less noise interference than that without processing (Fig. 3F).

After successfully extracting the edges of cells, the next step is to transform the line segments into definite boundary information, that is, the fitting process. Hough detection is a fitting algorithm, which maps the corresponding parameters of the line segments in the image into the parameter space, and then solves them in the parameter space. Hough transform operation is essentially a statistical voting process. Point information in Euclidean space is transformed into line segment information in parameter space, which can be visually regarded as line segment parameters in Euclidean space.

Hough transform has developed from the earliest line fitting, which gradually includes the detection of various basic graphics. The whole algorithm flow can be easily realized by OpenCV library function. As an important basic figure, circle detection has important application in computer image recognition. When using Hough algorithm to detect a circle, the coordinates of the center of the circle (x, y) and radius R are taken as unset parameters. The experimental results are summarized (Fig. 3G).

#### Development of Cell Extraction Program

Some functions and functions based on OpenCV:

Cv2. Canny (). It is used to detect the external edges in the graphics, connect the detected points to the detected edges through threshold setting and output them.

Cv2. Finding Contour (). It can be used to detect closed patterns in pictures. The input of the function is a binary image, and the output value is the closed pattern contours of the binary image. Therefore, the function can be used to complete the edge acquisition of the complete object in the picture.

Cv2. Hough Circles (). Hough circle detection, in addition to cells, there are other interference and noise in the image. The image after edge detection inevitably has the interference of non-target edge, so it is necessary to set the fitting parameters and the geometric size of the fitting pattern for further image segmentation and extraction. Hough detection method is used to remove the wrong circle detection caused by size mismatch. The test results are shown in the figure. Para is further adjusted to adjust the degree of fitting so as to eliminate possible over-fitting phenomena.

Through centralized processing of 20 photos, the parameters with better comprehensive extraction effect are obtained, and the 20 pictures were tested. 338 cases were identified manually and 305 cases were identified by machine. The accuracy rate of cell recognition was 90.2%. The parameter range is determined: Hough detection Para2 is set to about 40. In minRadius about 50, maxRadius about 100 is better.

Fixed parameter detection, by setting minRadius to 40, maxRadius to 90 to detect more than ten pictures of cells, the results are good. (Table 1)

**Table 1.** Detection of the first type of cell number.

| Image ID | Number of cells detected | Number of human eye recognition cells |
| --- | --- | --- |
| 1 | 20 | 19 |
| 2 | 31 | 32 |
| 3 | 26 | 25 |
| 4 | 26 | 23 |
| 5 | 74 | 68 |
| 6 | 83 | 75 |
| 7 | 74 | 72 |
| 8 | 82 | 79 |
| 9 | 67 | 65 |
| 10 | 80 | 77 |

#### Nuclear cytoplasm extraction

Nucleus plays an important role in cell recognition. We can see that some of the cells have good imaging effect. The color of the nucleus is clearly distinguished from other parts of the cell. We consider separating the nucleus by combining edge contour search with morphological changes. Because Canny arithmetic can only extract the outer edge of cells better. Therefore, Otsu segmentation is used. First, the cells are segmented by threshold. It is found that there is a layer of “white circle” around them, which is caused by the poor imaging of microscopy. Corrosion operation is a morphological operation. We need to set the size of convolution core and the number of iterations by ourselves. Then the convolution core slides along the image, and the values of all areas covered by the core are true (in the experiment, the white part of the cell). Then the central pixel of the core keeps the original pixel value, and vice versa, changes to the background. After operation, the pixels near the edge of the target will be set to the background, resulting in the target object will become smaller. Corrosion operations are used to remove noise in the background and to uncover partial adhesion between detection targets. Contrary to corrosion, the convolution core covers the pixel values in the original image window. When there was a 1, the central pixel is considered to be the foreground. Morphological operation is to corrode first and then expand, which makes the target size unchanged while removing small noise. On the contrary, closed operation is used to fill small black holes in the target. Fig.4 showed that the outer circle of Otsu results was an interference item, and there was also a little interference in the nucleus. Open-close operation could be used to solve the problem (Fig. 4A).

**Fig. 4.**
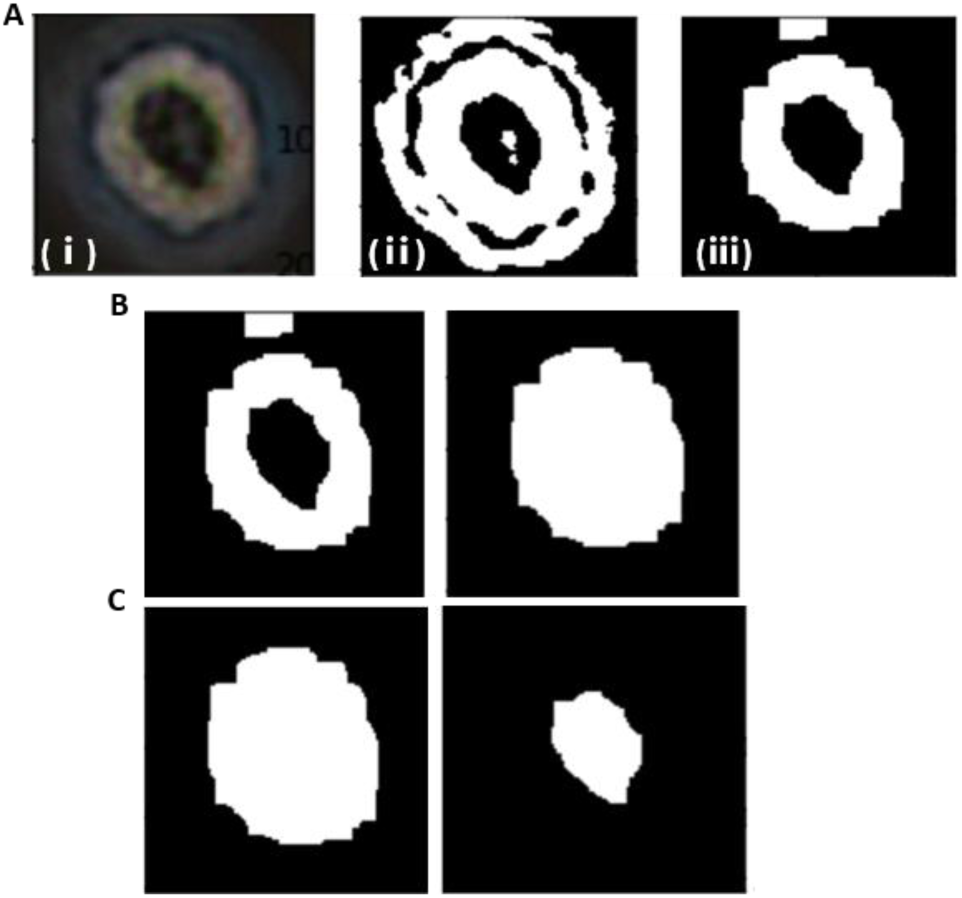
Nuclear cytoplasm extraction. (A) Cell binarization procession.i, Original image; ii, Otsu segmentation results; iii, Morphological manipulation. (B) Extracellular Contour Search. Left, Binary image; Right, Extracellular contour segmentation. (C) Extraction of nucleus. Left, total cell region; Right, Nuclear extraction.

Firstly, open operation is carried out to remove the interference of cell boundary, and clear cell boundary is obtained. Then closed operation is used to repair small loopholes. Further boundary searching is carried out, because the cell boundary in the searched contour is the largest, the maximum value is extracted by sorting the boundary, that is, the extracellular contour (Fig. 4B). Then, the black and white pixels of the processed image are reversed, filled, and then processed or calculated with the original binary image to get the core part (Fig. 4C).

In conclusion, the parameters of cell perimeter and area can be obtained approximately by using the fitting radius and perimeter of the circle. However, the problem of nucleus extraction process is that the method requires a high contrast between nucleus and cytoplasm. In the program design, the background brightness of cell images varies greatly because of the different light input, which affects the brightness of cells. In order to make the cells complete after the open-close operation, the setting of the open-close operation kernel function has a great change, which does not have universality.

#### The second kind of cell image: region extraction and watershed de-adhesion

At this time, the cells were segregated and partially adhered. Some of the cells adhered to the wall in a narrow and long shape, while others remained round (Figure 5B i). It is not feasible to use Hough changes, because the cells which account for the majority of the cells no longer have circular characteristics. OpenCV has findCounter () function, but it reads the binary graph, so it is necessary to transform the color map first, and the boundary information is not lost in the process of conversion.

**Fig. 5.**
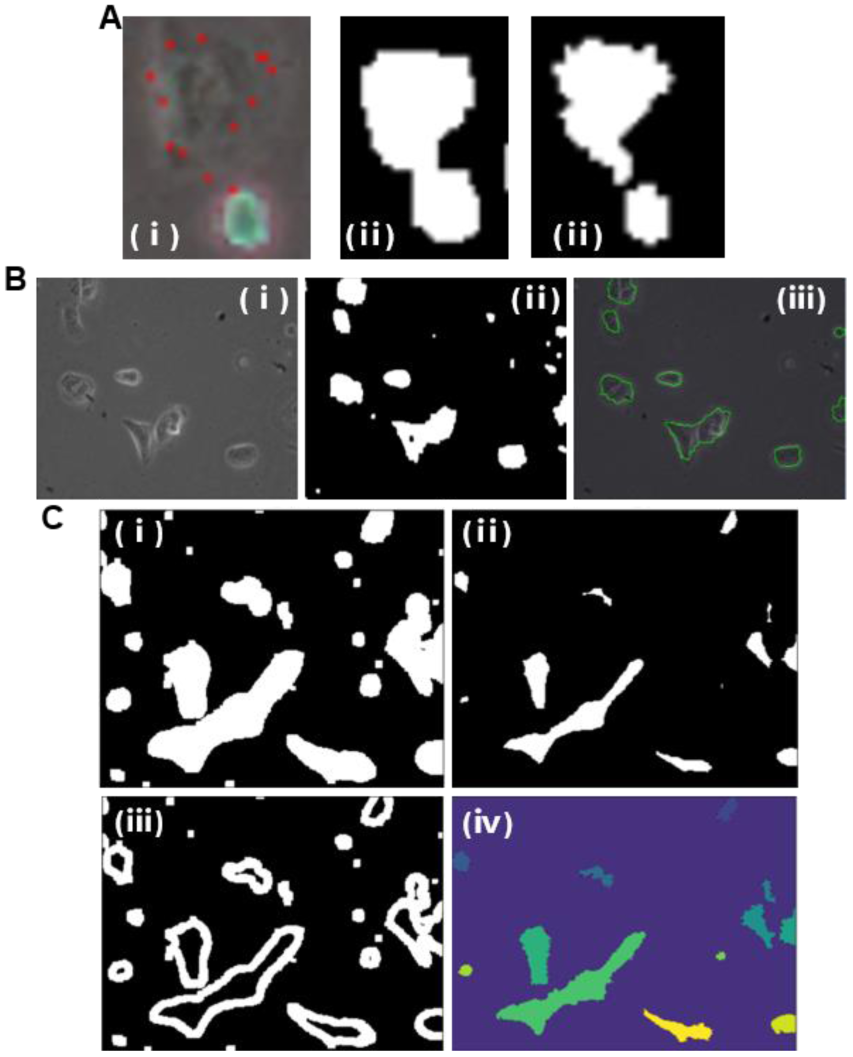
Segmentation and recognition of sparse adherent cells. threshold transformation combinedtraction of images; iv, The boundary and background of uncertain region are with watershed is considered for contour detection. (A) Original image, threshold segmentation and watershed treatment. (B) Original image, Binary image and segmentation results. (C) the procession of watershed algorithm for debonding. i, Expansion slightly enlarges cell boundaries; ii, Determining seeds by threshold segmentation after distance transform; iii, Obtaining uncertain areas by sub determined by image subtraction, and then watershed segmentation is performed.

First of all, it is necessary to complete the rough edge detection. We discovered that the binary threshold segmentation has great difference in brightness with the change of magnification and transmittance of the microscope, and it has low generality. While the effect of Otsu threshold method is better than that of binary segmentation (Fig. 5B ii). Otsu threshold method cannot get the complete edge after adjusting the kernel function of many open and close operations. When opening operations first, the boundary of darker cells will be filtered out, resulting in loss of information. When closing operations are carried out, noise points will be amplified to form binary patterns similar to smaller cells, which will bring difficulties to subsequent counting operations (Fig. 5A).

Canny detection can get a clear and comprehensive cell contour, but the disadvantage is that small noise is also used as a boundary, but after many experiments and setting a better threshold, the impurities become fine. When using smaller kernel function to open the operation, most of the noise can be filtered and the complete and undistorted cell image can be preserved.

After observing the cell images, it was found that the adherent areas of the cells after the above-mentioned operation had not been improved, and because the adherent channels of the closed operation part became larger, and the cell bank of the adherent part occurred more frequently in this stage, so the adherent part needed to be processed by technical means

Watershed segmentation is an image segmentation method. In the algorithm, the image is abstracted as a high-low scattered topographic map, where the height of each point is the gray value of this point. In this way, the individual center of the image will form a low-lying area, while the contact boundary of different individuals to be segmented will form a relatively high demarcation line.

#### The function of watershed in Opencv is: cv. watershed (image, markers)

In the practical application of the algorithm in this subject, the cell boundary is slightly expanded by expansion, and the region of the target in the segmentation is obtained (Fig. 5C i). Then the distance between each target pixel in binary image and its nearest zero-pixel point is calculated, that is, 0 is set for zero pixel, and for other pixels, the algorithm finds the geometric path to the nearest zero-pixel point around it, which is expressed by gray value. Then the seed points are determined by threshold segmentation (Fig. 5C ii). Then the uncertain region is obtained by image subtraction (Fig. 5C iii), and then the boundary and background are calibrated. Watershed segmentation is processing (Fig. 5C iv).

Next, we evaluate the algorithm and extract the parameters. The results of observation are as follows: The program can identify almost all cell boundaries; The lens stain will appear in the same area of each image before the algorithm is processed. It interferes with the imaging of cells and is improved after the whole process; Adhesion was partially improved (Table 2).

**Table 2.**
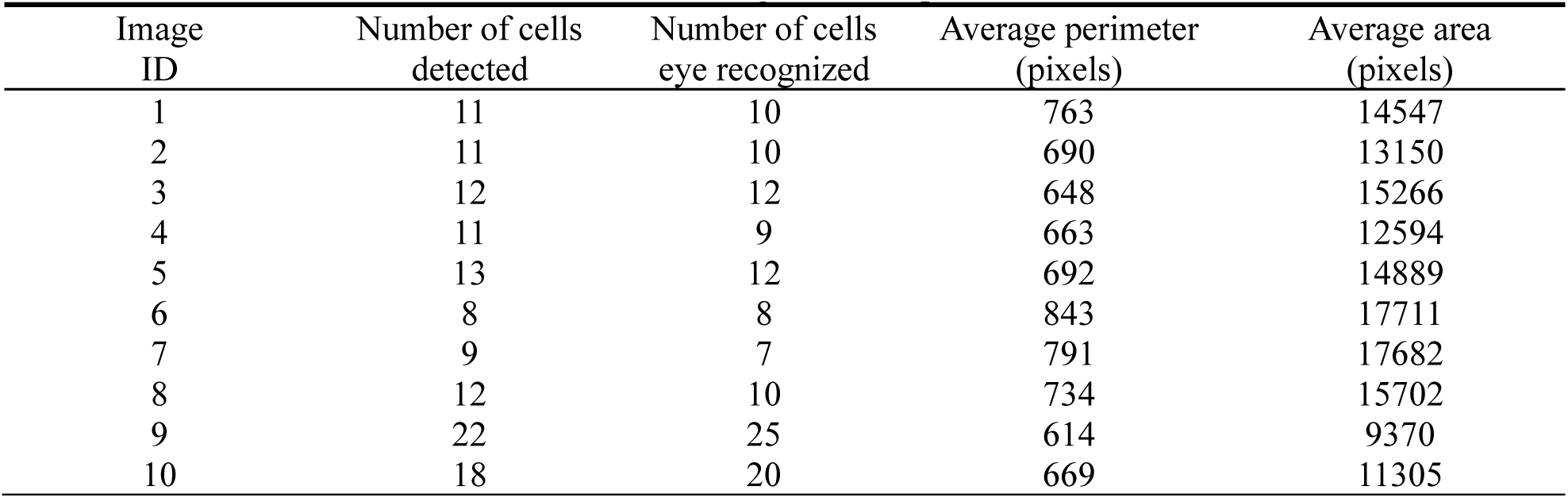
Extraction and Recognition of Sparse Adherent Cells.

Findcountour () and chain code were used to get the parameters of cell circumference and area. In the process of feature extraction, the functions of cv2. contourArea (), cv2. arcLength (), etc. of OpenCV can provide basic image information.

#### The third type of cell image: K-means Clustering Segmentation

These types of cells adhere densely to each other and have very small spacing. They can be divided into cellular and acellular regions. There is a certain color difference between the cell boundary and the cell in intercellular region. The cells showed irregular shape (Fig. 6A).

**Fig. 6.**
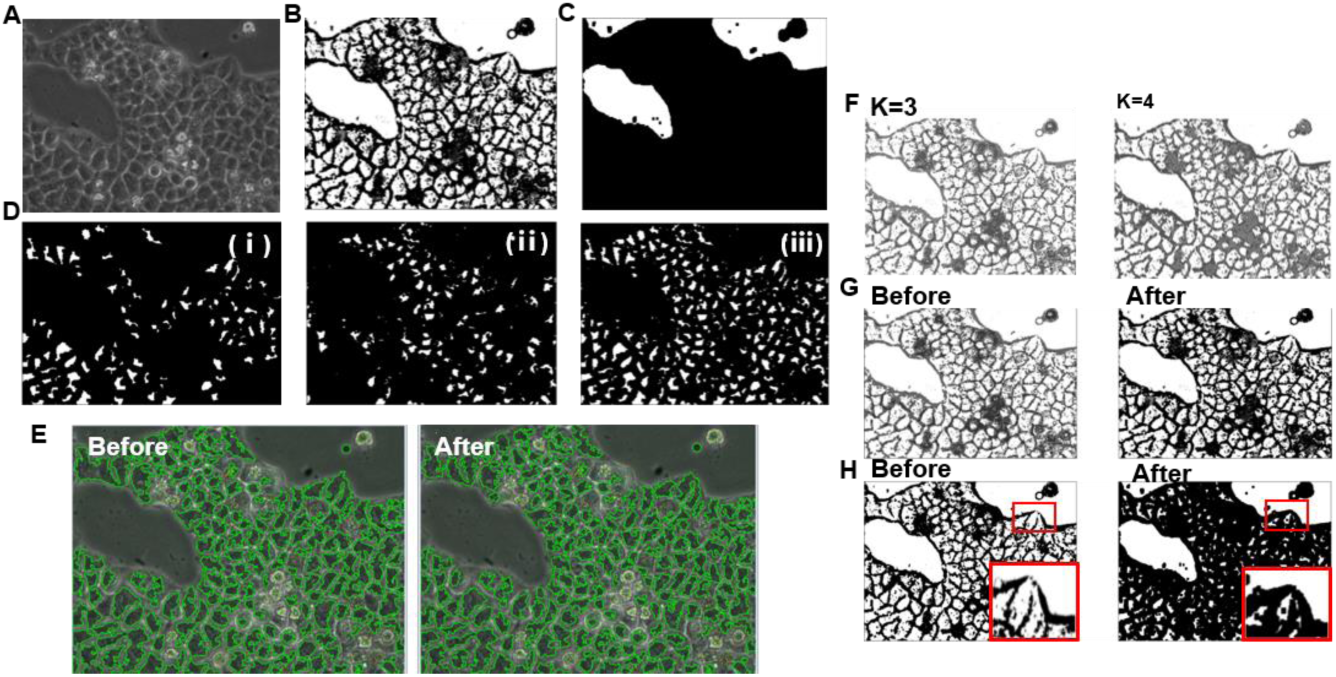
The processing of close adherent cell segment. (A) original image of close adherent cells. (B) image of mean clustering and image enhancement.(C) Image segmentation. (D) Segmentation result operation with different thresholds after distance change. i, Strong segmentation of larger cells. ii, Weak segmentation of small cells. iii, Consolidated results. (E) Comparison of watershed results with different thresholds after distance change. (F) K-means Clustering Results when k=3 and k=4. (G) Contrast before and after treatment of high cap and low cap. (H) Contrast before and after expansion operation.

K-Means algorithm is a classical unlabeled classification algorithm based on distance similarity. By comparing the distance between samples in multi-dimensional Euclidean space and adjusting the center point gradually, the samples are classified into different categories. When using K-means to classify colors, first of all, it is designated to be divided into several categories, that is, K value according to the actual situation. When the algorithm runs, the centers of each group are randomized at first, and then the distance from each sampling point to the centers of each group is calculated. The point is divided into the nearest group, and then the center position is updated averagely based on the position of each group. Repeat the following procedure until the center of each group remains unchanged.

The color of the central part of each adhesion cell is similar, and the color of the edge part is lighter. Therefore, K-means color classification can achieve better results. The color of each point in the color image is taken as the coordinate value in the multi-dimensional space. Image colors can be roughly divided into three or four categories for experiments (Fig. 6F). High HAT transform and low HAT transform can enhance image contrast, which we use the method to increase the chromatic aberration between cells for subsequent segmentation (Fig. 6G). After the processing, cells and large backgrounds are white prospects, which needs to be further separated by other treatments. And cell counting based on area statistics needs clearer and more accurate cell area. The total cell culture area is the key index to judge whether the cell is subcultured and proliferated. Therefore, it is necessary to extract the area of cell growth area and to distinguish the background and foreground accurately (Fig. 6C). We can see that the cells adhere to each other and join into large pieces. Consider using connected domain to distinguish them.

The judgment of connected area in OpenCV is based on the distribution of binary data of pictures. For binary images, if the binary values of adjacent pixels in the two fields are the same, then the two points are connected. From the image, the boundary of the cell region is black, which is obvious compared with other regions. Cell region can be regarded as a connected domain and other parts as another connected domain. Otsu method is used for threshold segmentation to complete the preparation of binarization. After binarization (Fig. 6B), in order to ensure that all boundaries are sealed, expansion treatment is carried out to complete the preparation of foreground and foreground (Fig. 6H).

After that, we can distinguish the cell block from the background precisely by setting the larger area as the background. It is noticed that some background is separated into several small pieces. Therefore, we should compare the area areas. For the larger white area, we can judge the background. Although the cell area is also white, it is larger after expansion. Large reduction, in area and background distinction is obvious. After morphological operations, the watershed algorithm of adherent cells was continued. It is noteworthy that in the process of experiment, when the threshold changes after the distance changes are noticed, if the increase of the threshold can better solve the adhesion, but the recognition of smaller cells is affected, small threshold segmentation is considered for smaller cells (Fig. 6D ii), larger threshold is set for larger cells (Fig. 6D i), and finally the results are merged (Fig. 6D iii). Fig. 6E showed the comparison of watershed results with different thresholds after distance change.

## IV Cell Classification and Recognition

After cell segmentation and extraction, the next work is mainly focused on cell recognition. We use the mature training model which has been developed and combined with the cell image which has been tailored and processed according to the specific input rules to train.

### Brush Selection and Design of Neural Network Model

BP neural network is a classical forward transfer network. The network structure includes input layer, hidden layer and output layer. The number of neurons and learning rate, etc. At the beginning of training, the weights and other parameters of each neuron in each layer are initialized randomly, and then the training is started. For each sample in the training set, firstly the network calculates the results forward according to the network structure of the current model, and then updates the neuron parameters and the corresponding bias of each layer in the network according to the deviation between the real value and the calculated value calibrated beforehand. Considering the area, perimeter and other characteristics mentioned above, it is possible for the establishment of BP network.

The first step is to extract the individual cell image from the image sample under the microscope; the second step is to extract the required features from these cell images by chain code; the third step is to train the model and get the weight file according to the obtained features. However, the performance of this method mainly depends on cell segmentation, image preprocessing, feature setting and other aspects of the algorithm design, especially the feature setting has great subjectivity. Therefore, it has great limitations. This paper uses convolutional neural network to extract features, avoids artificial feature extraction, which can make full use of all kinds of information in sample images and solve the shortcomings of traditional classification methods such as BP, clustering and so on.

CNN network is a kind of artificial neural network which has developed rapidly recently. The standard construction of CNN includes input layer, convolution layer, pooling layer, full connection layer, various classifiers and so on. On this basis, more complex models such as RCNN have been developed. CNN shares weights in the network. The design of this network structure reduces the number of weights and thus reduces the computational complexity. It can directly use pictures as input and automatically extract features.

### Development environment and construction of neutral network

Keras is a Python-based advanced neural network API. Its code is readable and the construction of the network is simple and easy to use. And it can run as a back end with TensorFlow. The development environment of this subject is Windows 10, keras 2.2.4, tensorflow 1.9.0, and the system hardware is Intel i7-6700HQ and Nvidia 960M GPU with GPU hardware acceleration. In the process of installation, we need to install CUDA and graphics card driver corresponding to the version.

Model.add (): Add layers to the network constructor layer by layer.

Model.compile(): Before the training model, the optimizer and evaluation criteria in the learning process are configured.

The input image size is W ×W, the filter size is F × F, and the step size is filled with P pixels. The output image size is N×N, where N=(W-F+2P)/S+1.

First, the label settings of the images are completed through Python. In the classification and selection of cells, different cells have different proliferation rate after adherence growth and different protein expression, which leads to different cell morphology. After adherence, nerve cells have synapses; fibroblasts are fusiform; some cancer cells are round; some cancer cells are fusiform, while pluripotent cells are aggregated. However, all cells are round after digestion into single cells, other cells have no obvious morphological difference, except for oocyte and other germ cells which are easy to distinguish because of size. Therefore, it is more valuable to identify adherent cells. As like DU145 cells and MCF-7 cells exhibit unique morphology after tight adhesion. So, we selected DU145 and MCF-7 for cell identification and screening (Fig. 7A). Firstly, the training set pictures are divided into two categories. The DU145 and MCF-7 cell pictures are represented by 0 and 1 respectively, and the labeling of the samples is completed.

**Fig.7.**
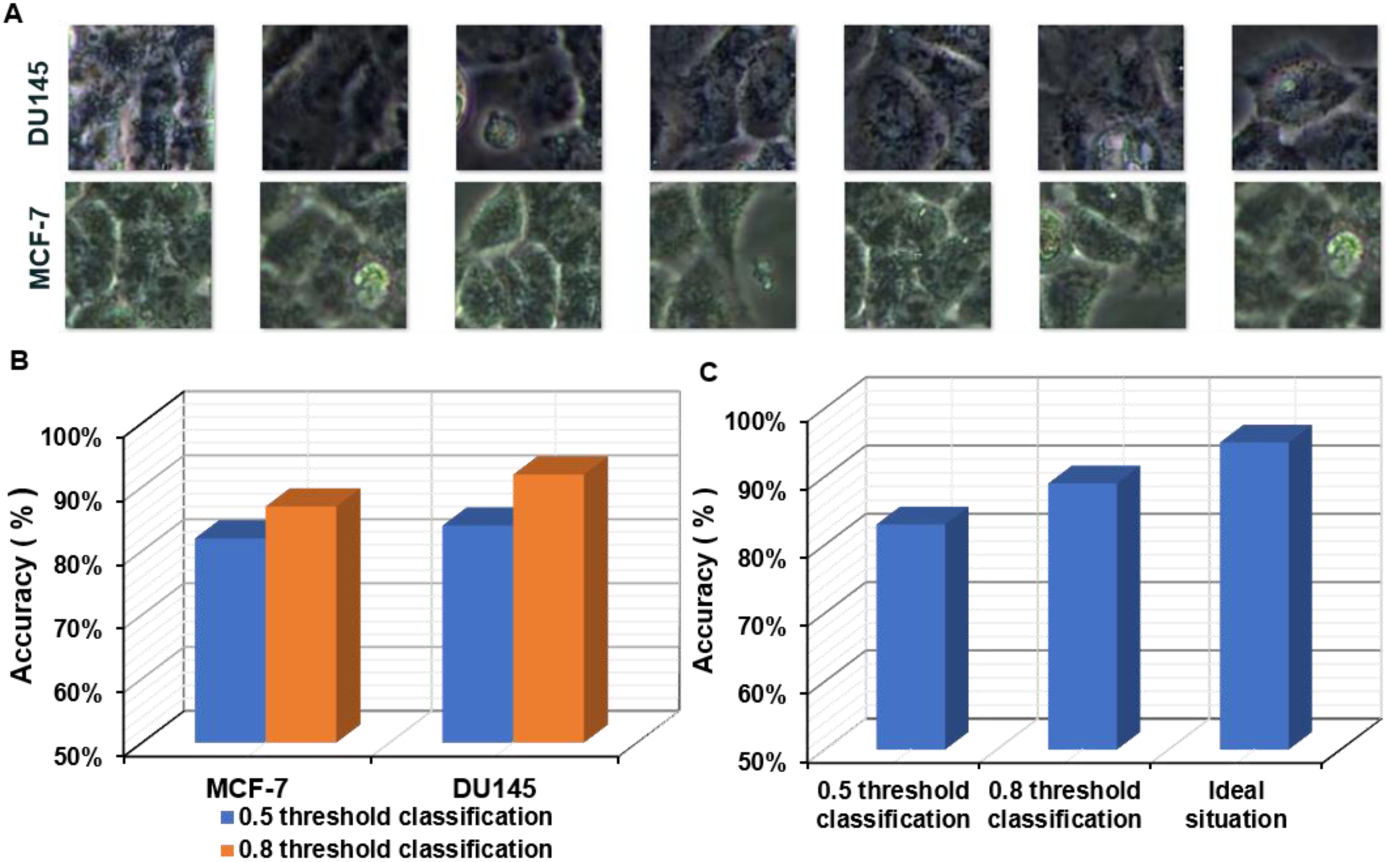
Cell classification and recognition. (A) Results of DU145 and MCF-7 cell taxonomy. (B) Accuracy of DU145 and MCF-7 cell classification and recognition. (C) Overall situation forecast.

Firstly, a VGG-like convolution neural network is constructed, which consists of four convolution layers, two pooling layers and full connection layers. After training, it is found that the accuracy of the network is low and the loss is large during the training process. In the permission of machine and time, we cannot get satisfactory results by adjusting parameters such as epoch and batch size many times. The accuracy is only about 0.5-0.6. We redesign the two-class network, which not only simplifies the network structure, but also greatly improves the accuracy and reduces the loss. In the training process of the neural network, many classical models set the size of the input picture to be equal in length and width, which will facilitate the design of the model. In order to ensure that the image is not affected by distortion, the original size of 2000 * 1500 is cut to 300 * 300 by clipping to facilitate input. In order to achieve higher recognition level, we use image enhancement to expand the capacity of training set by random rotation and other changes. It is found that accuracy fluctuates with the change of parameters. The more iterations the model has, the easier it is to get the results with high accuracy. Troughing comprehensively measure the computer’s computing power and then carrying out experiments, it is found that good results can be obtained when epoch is 70 times. Experiments results showed that the model works well and the accuracy of the test set is over 95 %.

After training, predict.py program was designed to re-examine 300*300 pictures. Set thresholds to 0.5 and 0.8. For example, when we get the result that the probability of MCF-7 is 0.6 when detecting MCF-7, For the former, the result is MCF-7, while for the latter, we can’t judge and do not take part in the correct and wrong statistics. The results are showed in supporting information (Table 3 and Table 4).

**Table 3.** The results of MCF-7 recognition.

| MCF-7 | Using 0.5 as threshold<br>dichotomy | Using 0.8 as threshold<br>dichotomy |
| --- | --- | --- |
| The correct cell<br>number identified | 41 | 35 |
| Error cell number | 9 | 5 |
| Accuracy | 82% | 87.5% |

**Table 4.** The results of DU145 recognition.

| DU145 | Using 0.5 as threshold<br>dichotomy | Using 0.8 as threshold<br>dichotomy |
| --- | --- | --- |
| The correct cell<br>number identified | 42 | 36 |
| Error cell number | 8 | 3 |
| Accuracy | 84% | 92.3% |

Fifty pictures of MCF-7 and DU145 were extracted from the pictures taken outside the training set and the test set according to the above method for testing. It can be seen that the accuracy rate is more than 80 % (Fig. 7B). When cells are cultured on a large scale, there are many pictures of cells, which can be trained on a larger scale and dichotomized with a higher threshold, and the accuracy rate can be higher. And in the previous training process, we can see that the model converges gradually and has high accuracy in the training process, which reflects the correctness of the model design, and we can Table 3 The results of MCF-7 recognition expect that it can get better training on larger data sets (Fig. 7C).

This chapter mainly carries on the cell screening. We designed a two-layer convolution and two-layer pooling convolution neural network. The original size of 2000*1500 image is 300*300 by random clipping to facilitate input, and the capacity of training set is expanded by image enhancement through rotation and other changes. Then the network is input and trained with pictures, the weight file is obtained and the prediction program is developed.

## Conclusion

With the rapid development of various technologies in machinery, computer and other fields, computer vision, machine learning, automatic control and other technologies are widely used in the field of biomedicine. It brings a good opportunity for large-scale cell culture and cell screening. In our research, computer vision technology is applied to the construction of cell bank, which improves the way of manual cell counting and improves the automation. At the same time, image segmentation and screening of cells were carried out by combining computer vision processing and machine learning, which provided a reference for large-scale cell bank construction and hospital automated pathological research.

Firstly, RAW264.7 and DU145 cells were cultured and subcultured for two to three times. After researching the characteristics of the image and brushing the existing image processing segmentation algorithm, we decided to use Canny detection and Hough detection to identify the circular and partially overlapping cells represented by RAW264.7. Meanwhile, median filtering was used in the cell pretreatment stage. After that, watershed algorithm is used to partially eliminate the adhesion of cells with partial adhesion, and then Kmeans clustering color segmentation with severe adhesion is completed. In order to solve the adhesion problem, an optimization algorithm was tried to extract the ratio of cell area to culture area accurately, which provided a reference for the automatic detection and culture of cell factory. On the basis of image segmentation, cell recognition is carried out. CNN was selected for cell recognition after comparison. At the end of this project, the algorithm of each part of the whole subject is optimized, and the experimental data are obtained. The experimental analysis proves the feasibility of the overall scheme.

There is still room for improvement in the current work. Pictures of cells are initially divided into three categories, and more elaborate designs are needed in the actual industrial cell culture process. In addition, the image segmentation of strong adhesion cells in the segmentation algorithm is not perfect, and if there are more pictures, it can support the training of more complex neural networks.

## Conflicts of interest

The authors declare no conflicts of interest.

## Acknowledgements

The authors gratefully acknowledge the financial support from the Natural Science Foundation of Beijing Municipality (Grant No. 17L20128) and Beihang University (No. ZG216S1751, ZG226S188D

